# Interaction between Model-based and Model-free Mechanisms in Motor Learning

**DOI:** 10.1101/2025.04.28.651014

**Authors:** Adith Deva Kumar S, Adarsh Kumar, Neeraj Kumar

## Abstract

Motor learning can be driven by distinct mechanisms—habitual, model-free processes, and strategic, model-based processes—depending on the magnitude and context of movement errors. Although small and large errors are known to engage distinct motor learning mechanisms—model-free (implicit) or model-based (explicit), respectively—it remains unclear whether successfully deploying one mechanism might hamper the engagement of the other in subsequent learning tasks. Here, we investigated how prior engagement of a particular mechanism biases future adaptations, even when the new task context typically favours the alternative strategy. Across three experiments (N=82), participants performed reaching movements to targets that either remained fixed or “jumped” mid-movement by small (15°) or large (30°, 45°, or 60°) angles. When first exposed to small errors (15°), participants exhibited persistent aftereffects in subsequent catch trials and stable reaction times (RTs), hallmarks of a model-free, habitual process. Surprisingly, even when switching to larger error magnitudes later, these participants continued to show robust aftereffects and did not elevate RTs— indicating a carryover of model-free learning. Conversely, participants who initially experienced large errors showed minimal aftereffects and flexible RT modulation consistent with model-based strategies; this bias persisted in later sessions with smaller errors, leading to reduced habitual aftereffects. Notably, inserting a washout phase to reset baseline performance did not abolish these mechanistic biases, highlighting that the initial engagement of either model-free or model-based processes leaves a durable imprint on subsequent adaptations. Taken together, these findings demonstrate that motor learning is shaped not only by ongoing task demands (e.g., error magnitude) but also by an individual’s prior learning history. Understanding how initial learning experiences constrain future adaptations has broad implications for designing interventions and training protocols in motor rehabilitation and skill acquisition.

**Statement of Significance:** Motor learning involves distinct mechanisms: habitual, model-free processes (driven by gradual stimulus-response associations) and strategic, model-based processes (guided by explicit adjustments). This study demonstrates that initial engagement of one mechanism biases subsequent adaptations, even when task demands shift to favor the alternative. The findings suggest that motor learning is a hierarchical process shaped by cumulative contextual experiences. Our results have highlighted how early learning establishes neural or cognitive frameworks that constrain future adaptations, prioritizing efficiency over flexibility. This has implications for designing motor training or rehabilitation protocols: initiating learning with model-based strategies (via large errors) may preserve adaptability, while model-free training (via small errors) risks anchoring rigid habits. By elucidating how prior mechanisms bias ongoing learning, this work advances our understanding of motor memory interactions and their real-world applications in skill acquisition and recovery.

## Introduction

One of the primary functions of our brain is to coordinate movements by integrating visual, proprioceptive, auditory, and other sensory inputs to produce coordinated actions by controlling the muscles. Motor learning, a critical component of this coordination, involves practice-dependent changes in motor performance (Krakauer et al., 2019; Wolpert et al., 2011). Motor adaptation, a type of error-based motor learning, occurs in response to changes in the environment or body, allowing individuals to recalibrate their movements to maintain performance levels (Kim et al., 2021; Morehead et al., 2017; Shadmehr et al., 2010).

Multiple mechanisms are thought to underlie motor adaptation. These mechanisms have often been categorized by differences in learning rates (fast versus slow processes), the nature of feedback signals (reinforcement versus error-based learning), or the degree of conscious strategy involved (explicit versus implicit processes) (Haith & Krakauer, 2013; Smith et al., 2006).These perspectives largely converge on two overarching classes: model-based processes, which rely on an internal representation of the perturbation (e.g., forward or state-space models; Cheng & Sabes, 2006), and model-free processes, in which stimulus–response associations are strengthened without constructing an explicit model (Sutton & Barto, 2018). Model-free learning is typically gradual and implicit, often leaving robust after-effects, whereas model-based learning is faster, strategic, and can be voluntarily suppressed (McDougle et al., 2015).

Motor adaptation is primarily driven by sensory prediction errors (SPEs)—the mismatches between predicted and actual sensory outcomes of movements (Franklin & Wolpert, 2011). Typical experimental paradigms investigating SPE-driven adaptation involve perturbing the visual or mechanical feedback during movements. Examples include rotating the visual cursor, altering visual feedback gain, or introducing mechanical loads to the limb (Shadmehr et al., 2010; Franklin & Wolpert, 2011). In these paradigms, SPEs directly inform adjustments in the motor system’s internal model to reduce future prediction errors.

In addition to SPEs, motor adaptation can also have contribution from task errors (TEs), which reflect failures to meet movement goals—such as missing a target altogether (Bond & Taylor, 2015; Tsay et al., 2021; Oza et al., 2024). Certain paradigms, notably target-jump tasks, isolate TEs by shifting the movement goal mid-reach without altering sensory feedback. Such goal perturbations induce adaptation purely through task errors, without generating sensory prediction errors (Tsay et al., 2021).

The magnitude of errors plays a crucial role in determining the type of learning mechanism engaged. Small errors, often below the threshold for conscious detection, typically recruit implicit, model-free learning mechanisms. These model-free mechanisms gradually reinforce stimulus–response (S–R) associations, leading to robust aftereffects and habitual responses (Morehead et al., 2017; H. E. Kim et al., 2018). Conversely, larger errors typically engage explicit, model-based learning strategies. Under such conditions, participants become consciously aware of performance errors and strategically adjust their movements, often deliberately aiming away from the original target location to compensate (Bond & Taylor, 2015; Leow et al., 2018; Huberdeau et al., 2015).

This distinction between model-based and model-free mechanisms is clearly evident in target-jump paradigms, where only task errors (and no SPEs) drive adaptation. Large target jumps (e.g., 45° or more) predominantly trigger cognitive strategies such as explicit mental rotations or conscious re-aiming toward shifted targets (McDougle et al., 2015). By contrast, smaller target jumps (e.g., ∼15°) typically induce implicit recalibration and produce persistent aftereffects, reflecting automatic S–R associations that are resistant to conscious suppression (Sadaphal et al., 2022). Model-free mechanisms, although computationally efficient, produce inflexible and automatic responses, whereas model-based strategies, despite their flexibility, require additional cognitive resources (Haith & Krakauer, 2013; Huberdeau et al., 2015). Thus, target-jump tasks provide a valuable approach to study how error magnitude dictates the balance between strategic flexibility and habitual automatization.

An important unresolved question is how these two learning mechanisms interact across sequential learning episodes, particularly when subsequent tasks demand a mechanism different from the one initially engaged. Recent evidence suggests that model-based and model-free processes are not entirely independent but instead interact and potentially compete for neural and computational resources during motor adaptation (Albert et al., 2022). Such competitive interactions could mean that initially engaging one mechanism influences how readily the other mechanism is recruited in future tasks. Thus, a critical open question is whether initial reliance on one mechanism—either habitual, model-free processes or strategic, model-based processes—biases the selection and effectiveness of the alternate mechanism in subsequent adaptation sessions. Clarifying this interaction would shed important light on how motor learning generalizes across contexts and could inform strategies to optimize training and rehabilitation approaches.

We hypothesize that initial engagement of either the model-free or model-based mechanism will bias subsequent adaptation, such that learners preferentially continue employing the previously successful mechanism even when a different mechanism would normally be optimal. We investigate this hypothesis using aftereffects and reaction times as indicators of model-free versus model-based processes, respectively. Our study directly addresses how learning a target-jump task through one mechanism impacts the selection and deployment of mechanisms in subsequent tasks, providing insight into the adaptive flexibility and generalization of motor behavior.

## Methods and Procedures

### Participants

A total of 82 healthy, right-handed individuals (mean age = 22.95 ± 3.37 years; 33 females) were recruited for the study. Handedness was assessed through Edinburgh Handedness Inventory (Oldfield, 1971). Participants had normal or corrected-to-normal vision and no history of neurological or orthopaedic disorders. All participants provided informed consent, were compensated monetarily for their time. All the experimental procedures and protocols were approved by the Institute Ethics Committee.

### Experimental setup

Participants performed point-to-point reaching tasks using a stylus on a digitizing tablet (GTCO CalComp). A HD monitor (1920 × 1080 Pixels) was mounted on top facing downward and projecting on a semi-transparent mirror placed in between the display and the tablet (Fig. 1A). Targets were projected from the display on to the mirror, creating a realistic depth view while occluding the hand. Cursor feedback, displayed as a yellow dot (0.3cm diameter) was provided throughout the movement, except during specific catch trials. Hand movement data were recorded at 120 Hz.

**Fig. 1.**
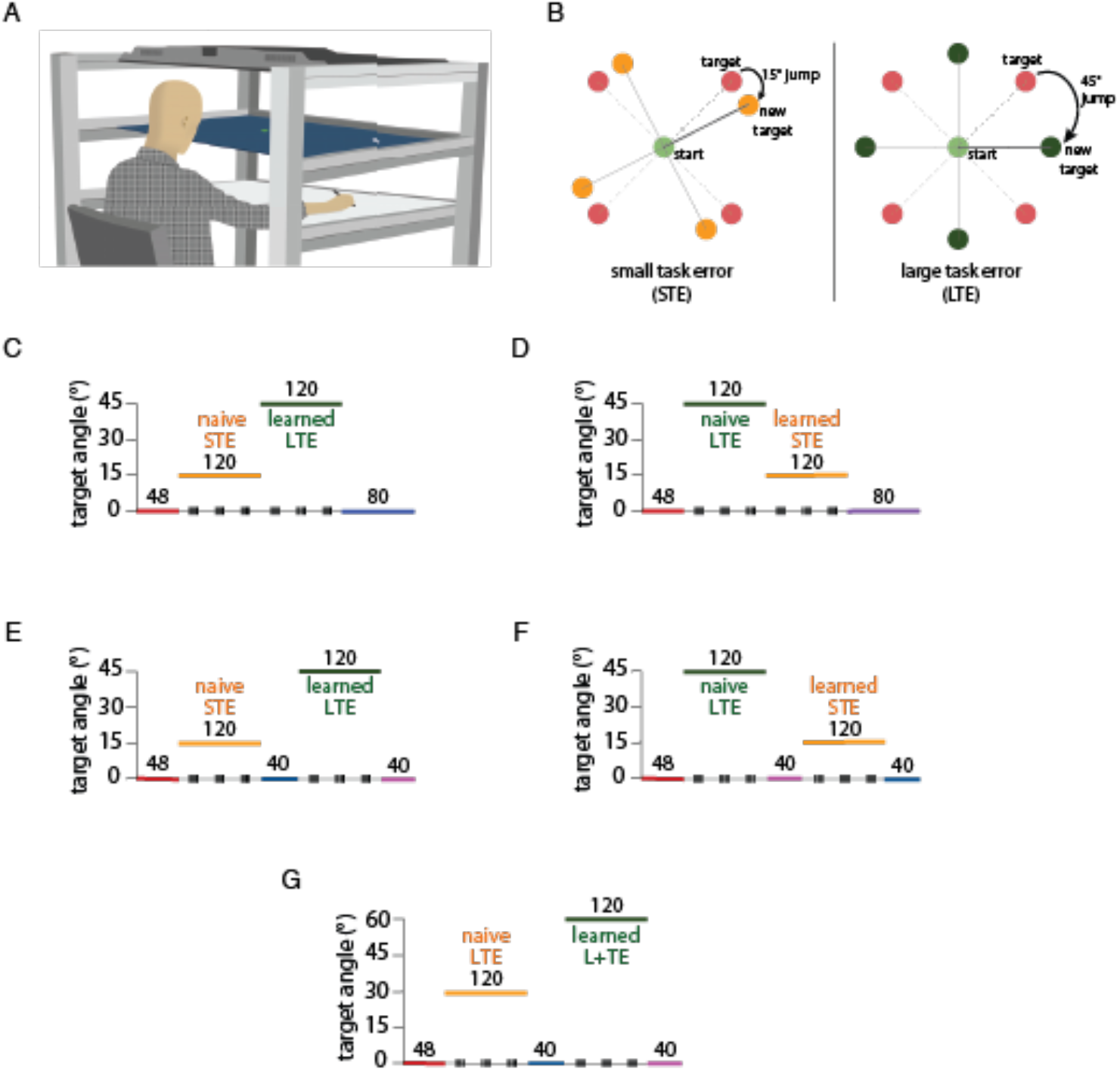
**A:** A VR experimental setup comprising an HD monitor mounted horizontally, projecting its display on the semi-silvered mirror placed at eye level, thereby obstructing the direct vision of the hand. A digitizer tablet placed below the mirror was used to make planar movements with a hand-held stylus. The mirror displayed the start point and the target along with the cursor, which represented the position of the stylus on the tablet, providing veridical feedback. **B:** Target location and jump locations in the point-to-point reaching task. A 15° jump (orange) induces a small task error (STE), primarily learned through a model-free mechanism, while a 45° jump (dark green) results in a larger task error (LTE), learned via a model-based mechanism. During the task, all targets were red circles, and the jump was implemented by removing the original target and instantly presenting the new target in the jump location. **C:** Task structure: subjects in the STE-LTE group, after a brief baseline session, were introduced to a small target jump of 15°, followed by a large target jump of 45°, and finally, a washout session. **D:** In the LTE-STE group, after baseline, large errors (45°) were introduced, followed by the small errors (15°) and the washout session. **E:** The STE-N-LTE group consisted of a baseline session, followed by a target jump of 15° and then a brief session of the washout session and the learning session 2 with a 45° target jump followed by the washout session again. **F:** The LTE-N-STE group performed a baseline session, a 45° jump session, a washout session, a 15° jump session, and a washout session. **G:** In the LTE-N-L+TE group, the baseline was followed by a large error of 30° followed by the washout, which was later followed by a much larger error of 60°, followed by the washout at the end. Three blocks of catch trials (four trials per block, one to each of the four targets) were interleaved in each learning session across all groups and experiments.

On each trial, participants were required to move the cursor to a start point (blue circle, 0.8 cm diameter) centered on the screen. They had to maintain this position for 500 ms, after which the start point turned green, signaling the appearance of a peripheral target (red circle, 0.8 cm diameter) at one of four pseudorandomized locations (45°, 135°, 225°, or 315°) 12 cm from the start point. Participants were instructed to reach the target within 1.5 seconds and hold their position until an auditory cue (beep) marked the trial’s end. Participants received a visual feedback about the accuracy of their movement through a numerical score (current trial score and cumulative score) presented on screen after each trial: 10 points for landing within the target, 5 points within 0.25 cm of its edge, 1 point within 0.4 cm, and 0 points beyond this range. The overall points were the summation of the points obtained during each trial. Participants were required to move with a velocity in range 35 cm/s to 50 cm/s and “Too slow” or “Too fast” in red was displayed on if they moved slower or faster than the valid range respectively. If they stayed in the required velocity range, “Good” in green color was displayed. The points obtained did not impact the compensation provided to the participants at the end of the experiment, and the points were not analyzed. Cursor feedback was provided during the entire movement and was veridical with the actual hand position.

Three distinct trial types were included in all experiments. Baseline trials involved reaching movements with targets remaining at their original displayed location and veridical cursor feedback provided throughout. Catch trials were characterized by no cursor feedback and no target jump, designed to assess residual aftereffects and model-free mechanisms. Jump trials featured a target jump to a new location immediately after the cursor crossed a 0.5 cm radius from the start point, with veridical cursor feedback maintained.

Experiments consisted of multiple distinct sessions. The baseline session included baseline trials to establish initial performance with accurate visual feedback. Learning sessions primarily consisted of jump trials to induce adaptation, interspersed with intermittent groups of four catch trials to measure model-free and assess participants’ ability to disengage strategies. Washout sessions involved catch trials conducted after learning sessions to evaluate the persistence of adaptation through residual aftereffects.

Participants received explicit verbal and visual instructions detailing the trial structure prior to each session type. For catch trials, participants were specifically instructed to discard any strategies employed during jump trials and aim directly at the original target.

Target jump magnitudes categorized the learning errors into Small Task Error (STE) for a 15° jump, and Large Task Error (LTE) for jumps of 30°, 45°, or 60°, depending on the experimental condition (Fig. 1B). Participants underwent 12 practice trials (8 baseline followed by 4 jump trials) for familiarization prior to the experimental sessions.

#### Experiment-1 (STE-LTE and LTE-STE)

In experiment 1 (N=32), subjects performed 48 baseline trials, followed by 120 trials of learning session I, 120 trials of learning session II, and 80 trials of washout session. Each of the learning sessions included the 12 interspersed catch trials and 108 trials of target jumps.

The target jump magnitude was constant across the learning session for all targets but the magnitude changes in the subsequent learning session. Participants were randomly divided into two groups which differed in the magnitude of the target jumps (small or large) learnt first followed by the other. The magnitude of the jumps was 15° for the small jump and 45° for the large jump (Fig. 1B) and participants either learned the small or large jump first followed by the other according to the group they belonged to (Fig. 1C, D).

#### Experiment 2 (STE-N-LTE and LTE-N-STE)

Experiment 2 (N=32), mirrored experiment 1, except that a washout session of 40 catch trials was presented between the two learning sessions to washout the learning from the first learning session. Similar to the previous experiment, here also, there were two groups of participants who either learned small target jumps (15°) in the first session and large jumps (45°) in the second (Fig. 1E) or vice-versa (Fig. 1F).

#### Experiment 3 (LTE-N-L^+^TE)

Experiment 3 was similar in design to experiment 2. This study involved a single group of participants (N=16) who learned a large target jump of 30° in the first Session and a larger target jump of 60° in the subsequent Session (Fig. 1G).

### Data pre-processing and statistical analysis

Hand movement data were analyzed using custom MATLAB scripts. Of the total trials, 0.52% were excluded due to stylus disconnection or failure to initiate movement. From the remaining trials, 2% were discarded during learning phases for extreme hand angles (±100°), and 0.91% of catch trials were excluded for angles exceeding ±30°. Hand position data were filtered with a 10 Hz low-pass Butterworth filter, and movement speed was derived by differentiating positional data. Movement onset was defined as the first timepoint where hand speed exceeded 5% of peak velocity. Reaction time (RT) was calculated as the interval between target appearance and movement onset. For baseline and catch trials, the hand angle were calculated as the angle between the line connecting the center of the start circle and the original target, and the line connecting the center of the start circle and the hand position at peak tangential velocity. For the target jump trials, vector from the new target was used to calculate the hand angle. Target jumps were counterbalanced across participants (clockwise for 50%, counterclockwise for 50%). Data from counterclockwise jumps were converted to clockwise by multiplying hand angles by −1. Hand angles were baseline-subtracted to remove directional biases. RTs were normalized to individual baseline RTs and were expressed as a percentage. Hand angle and RT data were binned by averaging across one cycle of four targets. We used estimation analysis (Ho et al., 2019) to perform the statistical tests since the effects are sensitive to the variability in the data and outliers. We report the confidence interval for the mean differences along with the legacy p-values. We used only the first catch trial from each sub-block to improve the robustness of effects and reduce the forgetting effect within the sub-block. We performed the same analysis on the average performance in four catch trials, and result show similar patterns.

## Results

Participants performed reaching movements to one of four targets under veridical cursor feedback (Fig. 1B). To dissociate learning mechanisms, we introduced target-jump trials of two magnitudes: Small Task Error (STE, 15°) and Large Task Error (LTE, 45°). Participants first learned to adapt to either a STE or LTE in an initial learning session followed by a subsequent session where the target jump magnitude was reversed: STE learners transitioned to LTE, while LTE learners transitioned to STE. This cross-over design allowed us to directly test how prior engagement of model-free (STE) or model-based (LTE) mechanisms biases subsequent adaptation to opposing error magnitudes. Participants were explicitly informed of the target jump and instructed to adapt by reaching to the new location. Three catch trials sub-blocks (4 trials each) were interspersed with each learning session, where participants were instructed to ignore target jumps and aim for the original target. This design allowed us to investigate whether initial learning establishes a dominant learning mechanism (e.g., habitual model-free vs. strategic model-based) that persists even when task demands change in the subsequent session.

### The mechanism engaged during initial learning persists in the subsequent session despite change in task demand

In experiment 1, participants in STE-LTE group (Fig. 2A) learned small target jumps (Small Task Error, STE, 15°) first, followed by a large target jump (Large Task Error, LTE, 45°). While in LTE-STE, participants learned large target jump followed by the small target jump (Fig. 2B). In both groups, participants adapted their hand movements effectively to target jumps of different magnitudes. By the end of first session, participants in naïve STE and LTE conditions learned to aim directly towards to the jumped location (mean ± SD = STE: 11.41 ± 2.81; LTE: 41.41 ± 7.31). However, participants learned these different magnitude of task errors by utilizing distinct learning mechanisms based on the error size.

**Fig. 2.**
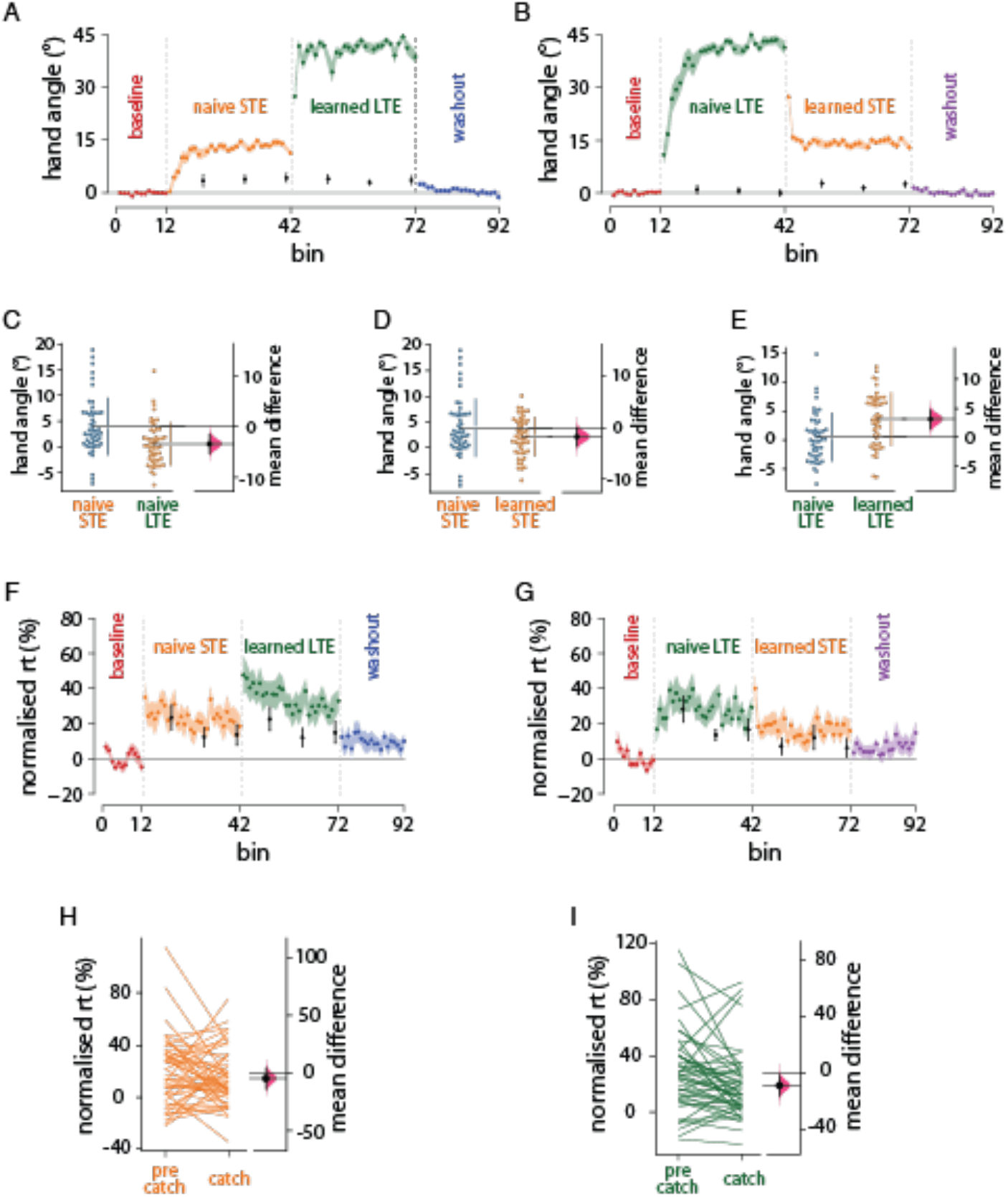
The mechanism engaged during initial learning persists in the subsequent session despite changes in task demand. **A:** The group averaged binned hand angle relative to the original target location in the STE-LTE group, showing participants adapting to the introduced jumps. Significant aftereffects were observed on catch trials during naive STE and persisted into session-2 despite the jump to LTE suggesting continuous persistence of model-free mechanism. **B:** The hand deviation in the LTE-STE group shows participants’ adaptation towards both jumps and the negligible aftereffects observed on catch trials in naive LTE get carried over to the later STE session indicating a more model-based mechanism throughout the sessions. **C:** Bootstrap t-test between the hand angle during the no-jump block of naive STE (STE-LTE group) and naive LTE (LTE-STE group) shows significant differences, suggesting the mechanisms used to learn these errors differ. **D:** Comparison between the catch trials of naive STE and learned STE shows that hand deviations during no-jump trials decreased when participants exposed to STE after learning through model-based mechanisms in the LTE condition. **E:** Naive LTE and learned LTE catch trials comparison shows increase in hand deviation in the LTE condition when participants had previously learned through model-free processes in the STE condition. **F & G:** Reaction time (averaged, binned and normalised) data of the STE-LTE and LTE-STE groups respectively shows decrease in RT over both learning sessions. **H:** In STE-LTE group, the naive STE reaction time remained stable across both jump and no-jump trials, suggesting inflexible, habitual stimulus-response execution which might not involve any strategy. **I:** In naive LTE, reaction time decreased significantly when transitioning from jump to no-jump trials, indicating rapid disengagement of explicit strategies.

To dissociate these mechanisms, we analyzed catch trials where participants were instructed to ignore jumps and asked to aim for the original target. In the naïve STE condition, significant hand deviations persisted during catch trials (mean ± SD = 3.95 ± 5.52, *t*_(50)_ = 5.10, *p* < 0.001, 95%CI = [2.39, 5.50], cohen’s *d* = 0.715), indicative of model-free stimulus-response (S-R) associations that resist voluntary suppression (Sadaphal et al., 2023). Conversely, negligible deviations in the naïve LTE condition (mean ± SD = 0.50 ± 4.13, *t*_(50)_ = 0.865, *p* = 0.391, 95%CI = [-0.661, 1.66], cohen’s *d* = 0.121) aligned with model-based strategic re-aiming, which participants could fully disengage when instructed (Taylor & Ivry, 2011). Critically, deviations in STE were significantly larger than in LTE (Fig. 2C, mean ± SD = STE: 3.94 ± 5.52; LTE: 0.50 ± 4.13, estimation statistics: [mean difference = -3.447, 95%CI = [-5.364, - 1.594], *p* < 0.001], [95%CI of cohen’s *d* = [-1.069, -0.280], *p* < 0.001] for two-sided permutation *t* test with 5000 bootstrap samples), confirming distinct mechanisms.

Prior work associates model-based learning with elevated RT (due to cognitive strategy formation) and model-free learning with RT similar to baseline (via automatized S-R mapping). Transitions from target-jump to catch trials revealed this mechanistic distinction. In naïve LTE, RT decreased significantly when transitioning from jump to catch trials (Fig. 2I, estimation statistics: paired mean difference = -8.932, 95%CI = [-16.423, -1.996], *p* = 0.017), reflecting rapid disengagement of explicit strategies. In naïve STE, RT remained stable across trial types (Fig. 2H, estimation statistics: paired mean difference = -4.750, 95%CI = [-14.001, 4.154], *p* = 0.311), consistent with inflexible, habitual S-R execution. This dissociation underscores that model-based learning allows adaptive strategy suppression, while model-free mechanisms engrain rigid responses.

These findings confirm that small task errors predominantly engage model-free processes characterized by persistent residuals of learned behavior and no change in RT when explicitly asked to disengage learned strategies, while large task errors invoke model-based strategies with adaptive disengagement of learned behavior.

We next investigated whether the mechanism engaged in the first session influenced learning in the subsequent session, despite changes in task error magnitude. In STE-LTE group, significant aftereffects were observed during naïve STE. These aftereffects persisted into Session 2 despite the transition to LTE, indicating continued reliance on the model-free mechanism. In LTE-STE group, negligible aftereffects during naïve LTE session were mirrored in the learned STE session, suggesting that the engagement of model-based mechanism persisted across sessions. We compared the hand deviations in naïve STE and learned STE during catch trials and found a significant decrease (Fig. 2D, estimation statistics: mean difference = -1.766, 95%CI = [-3.613, -0.014], *p* = 0.056) when participants were exposed to STE after learning through model-based mechanisms in LTE condition in comparison to naïve STE condition. Similarly, participants showed increase in hand deviation in catch trials in learned LTE condition after learning through model-free process in naïve STE condition (Fig. 2E, estimation statistics: mean difference = 3.094, 95%CI = [1.365, 4.679], *p* = 0.001). However, the contribution of model-free learning mechanism in the learned LTE session was not evident in the washout session. The hand deviation in the first bin of washout session after learning was similar (estimation statistics: mean difference = -1.078, 95%CI = [-2.724, 0.350], *p* = 0.192) in both groups. This could have been driven by the formation of deliberate strategies during later part of learned LTE session in the STE-LTE group.

Reaction time data showed similar bias in the second session (Fig. 2F, G). Participants could disengage the strategy during learned STE session and reduce the RT in catch trials (estimation statistics: paired mean difference = -8.153, 95%CI = [-14.515, -1.356], *p* = 0.015). We also observed similar behavior in the learned LTE condition as well that participants’ RT was decreased in catch trials from jump trials (estimation statistics: paired mean difference = - 14.225, 95%CI = [-25.442, -5.028], *p* = 0.006). This could have been driven by the upscaling of error magnitude and model-free system could not compensate the large errors.

These findings confirm the persistence of mechanistic biases across sessions—where prior engagement of model-free or model-based systems skewed subsequent adaptations— underscores how initial learning sculpts neural resource allocation. Participants defaulted to previously successful strategies regardless of new task demands, suggesting that early learning establishes a dominant computational framework that constrains future behavior.

### The mechanism carry-over is not due to an anterograde interference

The bias in learning mechanism during the second learning session could result from anterograde carryover of use-dependent changes in behavior. To test this, we designed the experiment 2, in which we introduced washout session after the initial learning of STE (Fig. 1E) or LTE (Fig.1F) where target did not jump, and participants were instructed to aim for the original target and disengage any strategy developed during learning. This also served to confirm whether participants engaged distinct strategies when learning small (STE, 15°) or large target errors (LTE, 45°).

Similar to experiment 1, participants learned to compensate both small and large task errors by aiming toward the jumped target location (Fig. 3A, B, [mean ± SD = STE: 10.92 ± 4.12; LTE: 43.45 ± 2.76]). Participants showed larger hand deviations in catch trials during naïve STE condition than naïve LTE condition (Fig. 3C, estimation statistics: [mean difference = -2.997, 95%CI = [-5.037, -1.066], *p* = 0.005], [95%CI of cohen’s *d* = [-0.954, -0.182], *p* = 0.005]). Further, during washout, the STE group exhibited significant residual hand deviations (estimation statistics: mean difference = -2.852, 95%CI = [-5.015, -1.533], *p* < 0.001), consistent with model-free S-R associations that resist disengagement. In contrast, the LTE group showed negligible aftereffects (estimation statistics: mean difference = -0.878, 95%CI = [-2.175, 0.099], *p* = 0.147), confirming that model-based strategies could be fully suppressed when instructed. Reaction time (RT) in the transition trials further validated this dissociation (Fig. 3F, G). In naïve LTE, participants rapidly reduced RT when transitioning from jump to catch trials (Fig. 3I, estimation statistics: paired mean difference = -22.396, 95%CI = [-33.032, -13.261], *p* < 0.001), consistent with adaptive strategy suppression. Conversely, RT remained stable across trial types in naïve STE (Fig. 3H, estimation statistics: paired mean difference = -1.517, 95%CI = [-9.436, 9.283], *p* = 0.760), underscoring the rigid, automatic nature of model-free learning. These results mirror Experiment 1 and validate that distinct mechanisms are engaged during small (model-free) versus large (model-based) error learning.

**Fig. 3.**
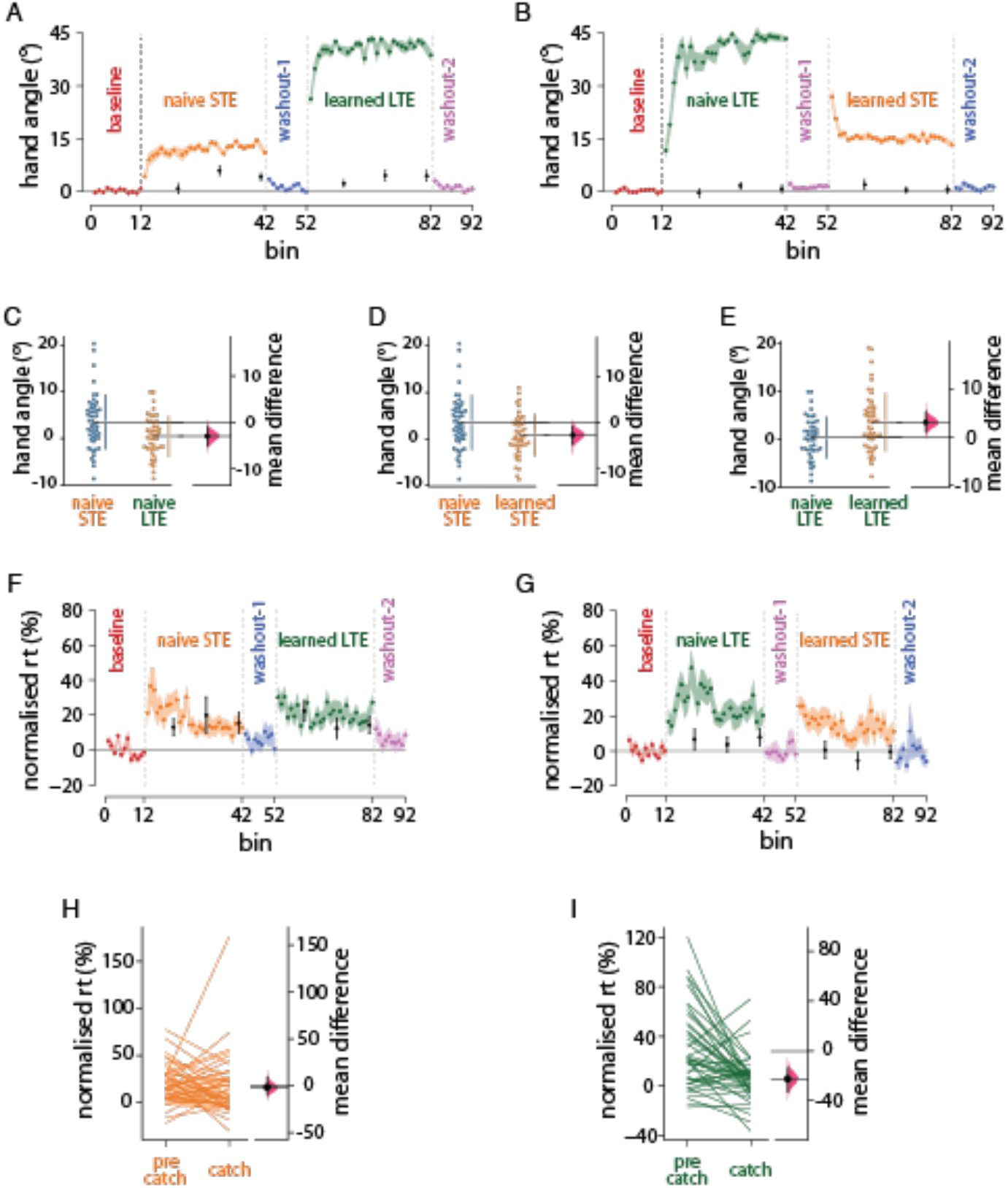
The mechanism carry-over is not due to an anterograde interference. **A:** In STE-N-LTE group, participants show visible learning towards both the jumps introduced and aftereffects were observed in catch trials of the naive STE session which is also observed in the learned LTE session despite learning being washed out in between. **B:** Evident adaptation was seen in both sessions in the LTE-N-STE group and the after effects were not present on the catch trials during the naive LTE as well as the subsequent STE session, even when washout trials were provided between them, These findings (A&B) suggest that the carry over of the mechanism is not due to anterograde interference. **C:** The hand angles on catch trials of naïve STE in STE-N-LTE group and naïve LTE in LTE-N-STE group differed significantly, implying that distinct learning processes were engaged. **D:** Catch trials of naive STE and learned STE comparison shows significant decrease in hand deviation when STE is learned after the model-based LTE. **E:** The learned LTE shows greater hand deviation on catch trials in comparison with naive LTE due to the persistence of prior learning of STE via a model-free mechanism. **F & G:** Normalised reaction time in the STE-N-LTE group and in LTE-N-STE group respectively. **H:** The reaction time during jump and catch trials of the naive STE session did not differ indicating the presence of rigid model-free mechanism during learning. **I:** Decrease in the reaction time during the catch trials in comparison with the jump trials in naive LTE reveals the involvement of the explicit model-based mechanism which can be disengaged on instruction.

After washout, participants transitioned to the second learning session (STE-N-LTE or LTE-N-STE). Strikingly, the mechanistic biases observed in Experiment 1 persisted. In the STE-N-LTE group, aftereffects on catch trials re-emerged during learned LTE session (mean ± SD = STE: 3.46 ± 5.77; LTE: 3.59 ± 5.92), with deviations resembling naïve STE (estimation statistics: paired mean difference = 0.127, 95%CI = [-1.926, 1.927], *p* = 0.904), suggesting latent model-free biases survived washout. Conversely, the LTE-N-STE group exhibited minimal aftereffects during learned STE session (mean ± SD = LTE: 0.46 ± 4.25; STE: 0.77 ± 4.46, estimation statistics: paired mean difference = 0.311, 95%CI = [-1.379, 2.185], *p* = 0.74), aligning with retained model-based flexibility. Direct comparisons revealed that hand deviations in learned STE were significantly smaller than in naïve STE (Fig. 3D, estimation statistics: mean difference = -2.686, 95%CI = [-4.777, -0.745], *p* = 0.011), demonstrating ability to disengage acquired strategies. Similarly, deviations in learned LTE exceeded those in naïve LTE (Fig. 3E, estimation statistics: mean difference = 3.124, 95%CI = [1.193, 5.162], *p* = 0.004), highlighting habitual carryover. Critically, this persistence of model-free behavior in the learned LTE session was evident even after resetting the S-R associations to baseline at the end of washout session (estimation statistics: mean difference = 0.254, 95%CI = [-0.583, 1.181], *p* = 0.588). Further, the hand deviations in the first bin of last washout session after learned LTE were larger than those observed after learned STE session (estimation statistics: mean difference = -1.973, 95%CI = [-4.288, -0.223], *p* = 0.076).

Reaction time data corroborate the persistence of initial learning mechanism in the subsequent session. Participants were able to disengage the strategy during learned STE session and reduce the RT in catch trials (estimation statistics: paired mean difference = -12.767, 95%CI = [- 17.884, -7.874], *p* < 0.001). Conversely, participants maintained the similar RT during catch trials as the learning trials in the learned LTE session (estimation statistics: paired mean difference = -1.318, 95%CI = [-8.430, 6.453], *p* = 0.726). This suggests that RT was also flexibly modulated based on the learning mechanism engaged during initial learning.

Experiment 2 confirms that the mechanistic biases observed in motor adaptation are not merely transient but reflect persistent computational preferences shaped by initial learning. Critically, mechanistic carryover persisted even after washout: prior engagement of model-free processes biased subsequent learning toward habitual S-R mapping, while model-based training promoted strategic flexibility in the subsequent learning session.

### Carryover of successful learning mechanism is not driven by scaling of error magnitude

One could argue that persistence of residual learning even in LTE session when learned after STE could be driven purely by the change in the task error magnitude instead of contribution of model-free processes. To test this, in experiment 3 (Fig. 4A, B), we investigated whether prior learning of a large task error (30°, LTE) will influence subsequent adaptation to even larger task error (60°, L+TE). Both errors fall within the large task error (LTE) range, which has previously been shown to engage model-based mechanisms.

**Fig. 4.**
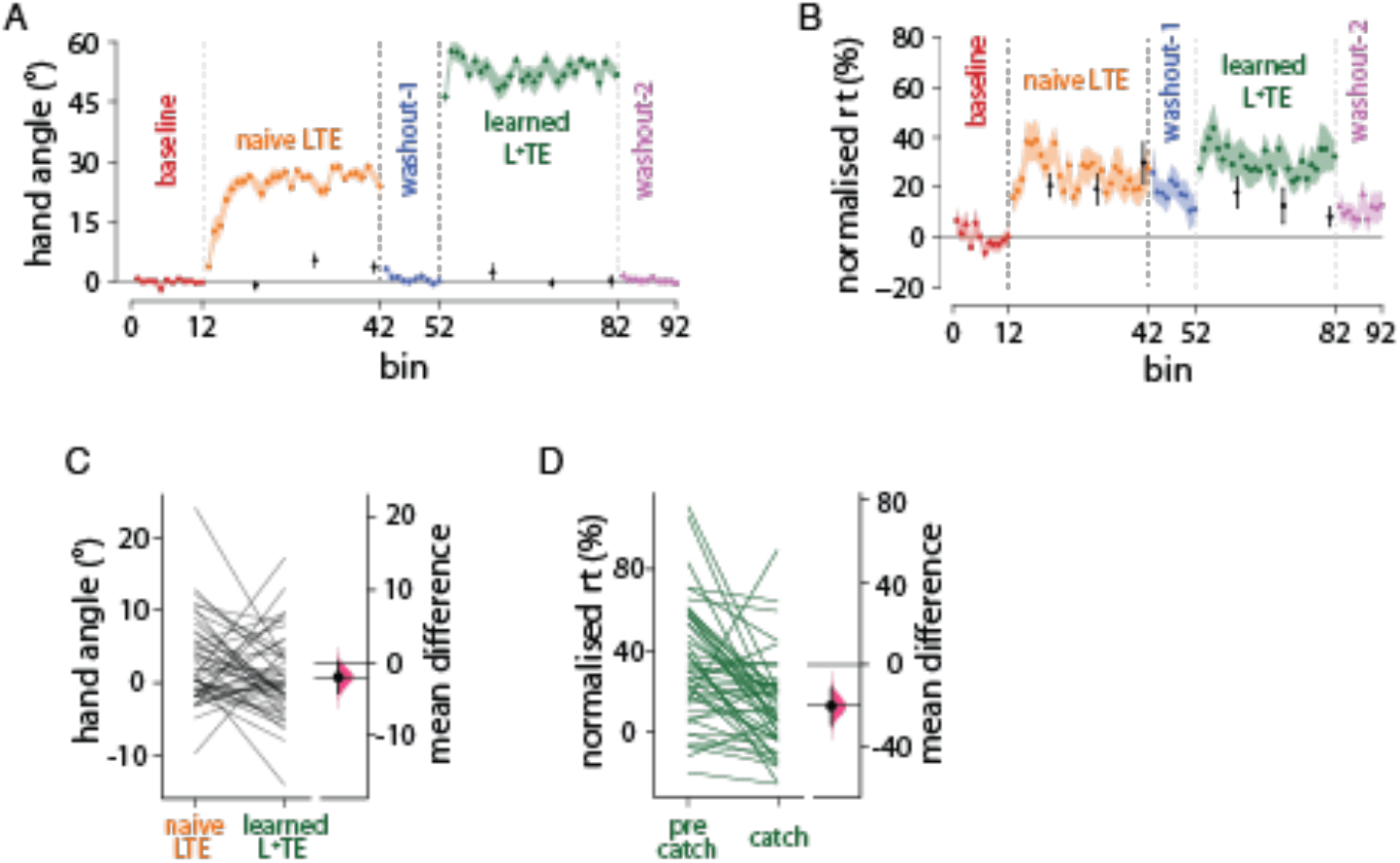
Carryover of successful learning mechanism is not driven by scaling of error magnitude. **A:** Hand movement angles during LTE and L^+^TE sessions show significant adaptation of participants to both the errors. **B:** The normalised reaction time during the sessions in LTE-N-L^+^TE group. **C:** The hand deviation on catch trials during naive LTE and L^+^TE sessions show no significant difference suggesting usage of same mechanism in both sessions. **D:** The reaction time on catch trials were evidently lesser compared to the jump trials during the L^+^TE learning session indicating the disengagement of the explicit strategy.

Catch trials revealed that hand deviation during L+TE (60°) were not significantly different from those observed in naïve LTE (30°) (Fig. 4C, estimation statistics: paired mean difference = -1.956, 95%CI = [-4.222, 0.253], *p* = 0.099). Similarly, participants were able to flexibly reduce the RT in catch trials from learning trials (Fig. 4D, estimation statistics: paired mean difference = -20.392, 95%CI = [-29.459, -11.641], p < 0.001]), indicating the disengagement of strategy in contrast to that observed in the learned LTE condition of STE-N-LTE group. This result suggests that participants employed a similar model-based strategy for both error magnitudes, and the learning mechanism did not require substantial adjustment when transitioning from 30° to 60°.

## Discussions

We investigated the interaction between model-based and model free processes during motor adaption to target jump task in reaching movements. Through a series of experiments, we show that, first, while small errors engages model-free, implicit learning mechanism which builds stimulus-response; large errors engage model-based explicit mechanisms. Second, the engagement of these learning mechanisms is limited by prior experience/use of the mechanisms in learning similar task. In experiment 1, we found that: Small errors (15°) predominantly engage model-free S-R associations, producing robust aftereffects and relatively unchanged RTs. Large errors (≥30°) recruit model-based strategies, yielding minimal aftereffects and RT decreases when switching from jump to catch trials. The mechanism engaged in an initial session persists in a subsequent session, even when the new session’s error magnitude typically favors the other mechanism. In experiment 2: we found that this bias remains even after a washout block to get rid of any anterograde effects. Furthermore, in experiment 3, when both errors are large (30° → 60°), participants simply reuse a model-based strategy without needing to shift to a new mechanism.

### Interaction between Model-Based and Model-Free Mechanisms

Our study examined how model-based and model-free processes interact during motor adaptation to target jumps in reaching tasks. We found that the initially engaged mechanism tends to persist across learning sessions, implying that once a learning mechanism is employed, it may continue to influence subsequent tasks, even if conditions change. Traditional views have held that these mechanisms operate separately, as seen in Mazzoni and Krakauer (2006), where implicit learning to correct sensory prediction errors could override explicit, strategy-based learning to reduce task errors. However, recent work (Albert et al., 2022) indicates that these processes can also interact and compete for resources, suggesting a more integrated motor learning system. Our findings align with this nuanced perspective, where an initial mechanism can influence later tasks, especially when prior experience has made that mechanism effective. This flexible interaction may enhance learning efficiency, potentially conserving cognitive resources by reusing successful learning pathways. The faster learning in subsequent sessions across our experiments supports this idea, indicating that if an initial mechanism (model-free or model-based) has been effective, the motor system “defaults” to it even under conditions that would normally recruit the alternate mechanism.

### Role of Error Magnitude in Mechanism Selection

Our findings emphasize that error magnitude plays a crucial role in determining the type of learning mechanism engaged. Small target errors (STE) predominantly activated model-free processes, while large target errors (LTE) engaged model-based strategies. This differentiation supports existing theories that error size influences whether motor adaptation is handled through habitual adjustments or explicit, strategy-driven recalibration. Large errors would have greater chances of inducing awareness about presence of error, leading to active engagement and disengagement of strategic, explicit cognitive neural processes (Sadaphal et al., 2022; Werner et al., 2015). In a recent study, Sadaphal et al. (2022) demonstrated that learning from small errors resulted in significant aftereffects in catch trials, reflecting the contribution of model-free processes that remain active without feedback. Conversely, for larger errors, participants used model-based mechanisms, with minimal aftereffects, supporting the idea that explicit strategies are engaged as needed and can be turned off when they are not required. However, our findings highlight that initial learning experience can override this typical assignment. Furthermore, Our Experiment 3 findings showed that adapting to a larger error (60°) after a smaller one (30°) was rapid, indicating that model-based processes allow scalable adaptation. This scalability enables flexible responses to increased error magnitudes without requiring relearning, contrasting with the relatively fixed nature of model-free associations.

### Persistence of Learning Mechanisms and Task Demand

The observed carryover effect across sessions highlights the adaptability of motor learning mechanisms in response to task demands. Our results suggest that, rather than being rigidly assigned to specific error types, model-free and model-based mechanisms adjust based on error size, task structure, and prior experience. Past studies often described model-free learning as rigid and resistant to modification based on task context. However, our findings align with the COIN model proposed by Heald and Wolpert (2021), which emphasizes that motor learning is context-sensitive, guided by cues from both present conditions and prior experiences. This perspective is supported by our results, which indicate that the mechanism selected can be influenced not only by the task’s immediate requirements but also by the learning history. This flexibility could allow individuals to optimize motor performance across varying tasks by dynamically adjusting which mechanisms are active based on both task demand and accumulated experience.

Interestingly, our Experiment 2 results showed that inserting a washout block, intended to ‘reset’ adaptation, did not diminish the mechanistic carryover effect; in some cases, it seemed to reinforce it. Washout trials are typically assumed to remove or weaken existing motor memories (Shadmehr & Brashers-Krug, 1997; Smith et al., 2006), the persistent bias we observed is strange at first sight. One possible explanation is that repeated exposure to the baseline condition after initial adaptation actually promotes consolidation of the earlier-acquired mechanism (Kim et al., 2018). The washout block may act as a distinct context or learning phase that, ironically, highlights rather than eliminates the previously successful strategy or S-R mapping. In line with the COIN model (Heald et al., 2021), the system integrates these contextual cues with prior experience, defaulting again to the initially successful mechanism when the subsequent task resembles the earlier perturbation. Thus, rather than erasing the carryover, the washout period may solidify it by helping the nervous system ‘tag’ the memory for future retrieval under related conditions.

Reaction time (RT) consistently differentiated model free vs. model based learning. In naïve conditions, introduction of large errors often prolonged RTs, reflecting the cognitive load of generating and updating an model-based strategy (Fernandez-Ruiz et al., 2011; Haith et al., 2015). By contrast, small-error adaptation did not alter RTs substantially, consistent with habitual or automatic stimulus-response (S-R) mapping characteristic of model-free learning. When switching from jump to catch trials, participants who had been using a model-based strategy rapidly reduced their RT (reflecting quick disengagement of explicit aiming), whereas those relying on model-free processes maintained stable RTs (reflecting an inflexible, habitual response).

Crucially, RT data also supported that prior learning experience biased subsequent adaptations. Participants who first learned with small errors (STE) tended to preserve lower or more stable RTs even when adapting later to large errors (LTE). In other words, they did not exhibit the typical elevated RTs usually associated with naïve large-error learning, suggesting that the model-free mechanism engaged during the first session continued to influence behavior. Conversely, participants who initially learned large errors (LTE) showed a sustained capacity for rapid RT modulation in the second session, even when the new error magnitude (STE) would normally favour a low-RT, model-free process. Here, the model-based strategy—once established—remained accessible and could be flexibly disengaged or scaled as needed, again reflecting a carryover from the initial learning phase.

### Neural Basis of the Interaction between Model-free and Model-based mechanisms

The persistence of the initial learning mechanism can be understood in the context of neural plasticity within cortico-striatal circuits. Model-based learning is thought to be mediated by the dorsomedial striatum and prefrontal cortical areas, which are responsible for flexible planning and updating internal models (Balleine & O’Doherty, 2010). In contrast, model-free learning relies on the dorsolateral striatum, a region implicated in habitual behavior and automatic S–R mapping (Tricomi et al., 2009; Yin & Knowlton, 2006). Our behavioral data suggest that initial exposure to small errors may strengthen the connectivity and plasticity of neural pathways associated with the dorsolateral striatum, thereby biasing subsequent adaptation toward a model-free framework even in contexts where a more flexible, model-based approach might be optimal. Similarly, explicit strategy formation during large error conditions likely reinforces neural circuits that support rapid adjustment and cognitive flexibility, which then persist even when error magnitude is reduced.

Electrophysiological studies provide converging evidence for this bidirectional interplay. Functional magnetic resonance imaging (fMRI) studies have shown that during decision-making tasks, activity in the dorsolateral striatum can bias the valuation signals in the dorsomedial striatum (Daw et al., 2011). Similarly, electrophysiological recordings using TMS-evoked potentials (TEPs) have demonstrated that as a behavior becomes more habitual, enhanced responses in the dorsolateral striatum modulate the sensitivity of the dorsomedial system when flexible adjustments are required (Jog et al., 1999). Behavioral experiments further support this reciprocal influence by revealing that strong habitual responses can constrain the adaptability of the cognitive map, occasionally resulting in perseverative errors when environmental conditions change (Yin & Knowlton, 2006). Conversely, when the two systems are well integrated, the model-free mechanism provides a robust scaffold upon which the model-based system refines decision-making, leading to more efficient adaptations (Daw et al., 2011).

### Summary

We asked how initial exposure to either small or large target errors, each recruiting different motor learning mechanisms (model-free vs. model-based), influences subsequent adaptations even when new error conditions typically favor the alternate mechanism. Small errors led to strong, habit-like aftereffects (model-free), while large errors triggered strategic re-aiming (model-based). Crucially, whichever mechanism was engaged first tended to persist in subsequent adaptation sessions, regardless of the current error magnitude. This mechanistic “carryover” remained intact even after washout periods intended to reset baseline performance. These findings reveal that initial learning experiences can bias future motor adaptations, suggesting that once a particular learning mechanism is established, the motor system “defaults” to it even under conditions that would ordinarily invoke the other mechanism. This has broad significance for optimizing rehabilitation and skill training, where the history of previous motor experiences could shape ongoing and subsequent learning.

## ACKNOWLEDGEMENT

We would like to thank IIT Hyderabad for all the institutional level support.

## Notes

CONFLICT OF INTEREST: None

FUNDING: This work was supported by DST CSRI (DST/CSRI/2021/164(C)(G)) grant to NK.

### Competing Interest Statement

The authors have declared no competing interest.

